## Supplement file for "Visual experience shapes functional connectivity between occipital and non-visual networks"

**This supplementary file includes:**

1. Figure S1 to S12
2. Table S1
3. Supplementary results

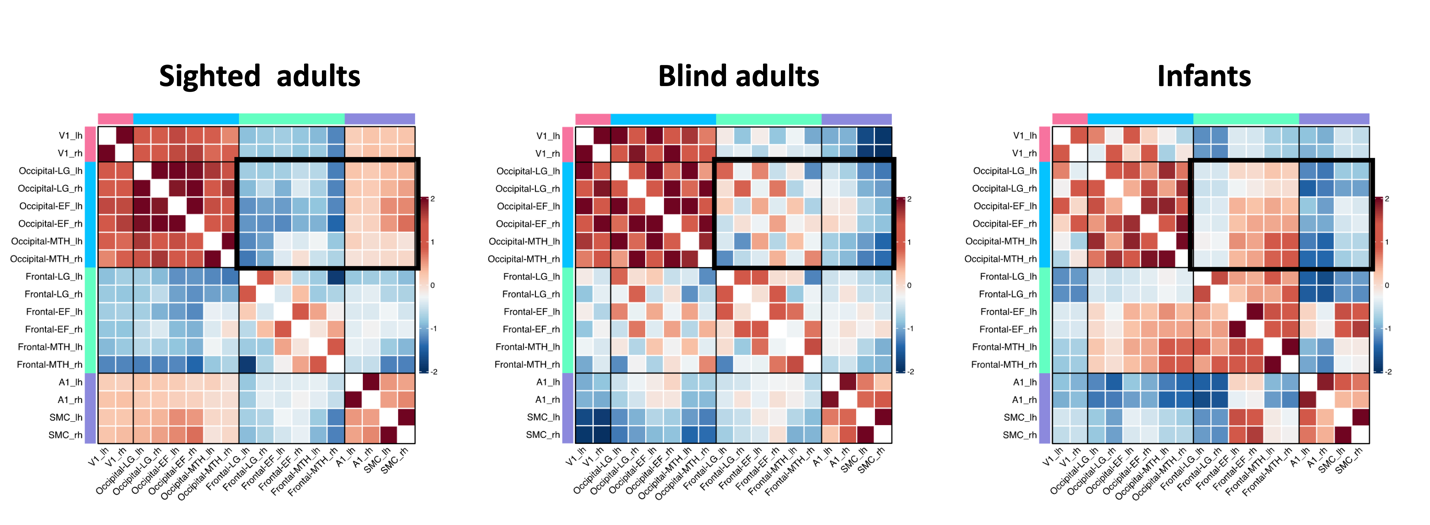

**Figure S1 The** **resting-state functional connectivity matrices in sighted adults, blind adults, and sighted infants.** The resting-state functional connectivity were normalized to ensure comparability across different groups. MTH: math-responsive region; LG: language-responsive region; EF: executive function (response-conflict) region; SMC: primary somatosensory and motor cortex, lh: left hemisphere; rh: right hemisphere.

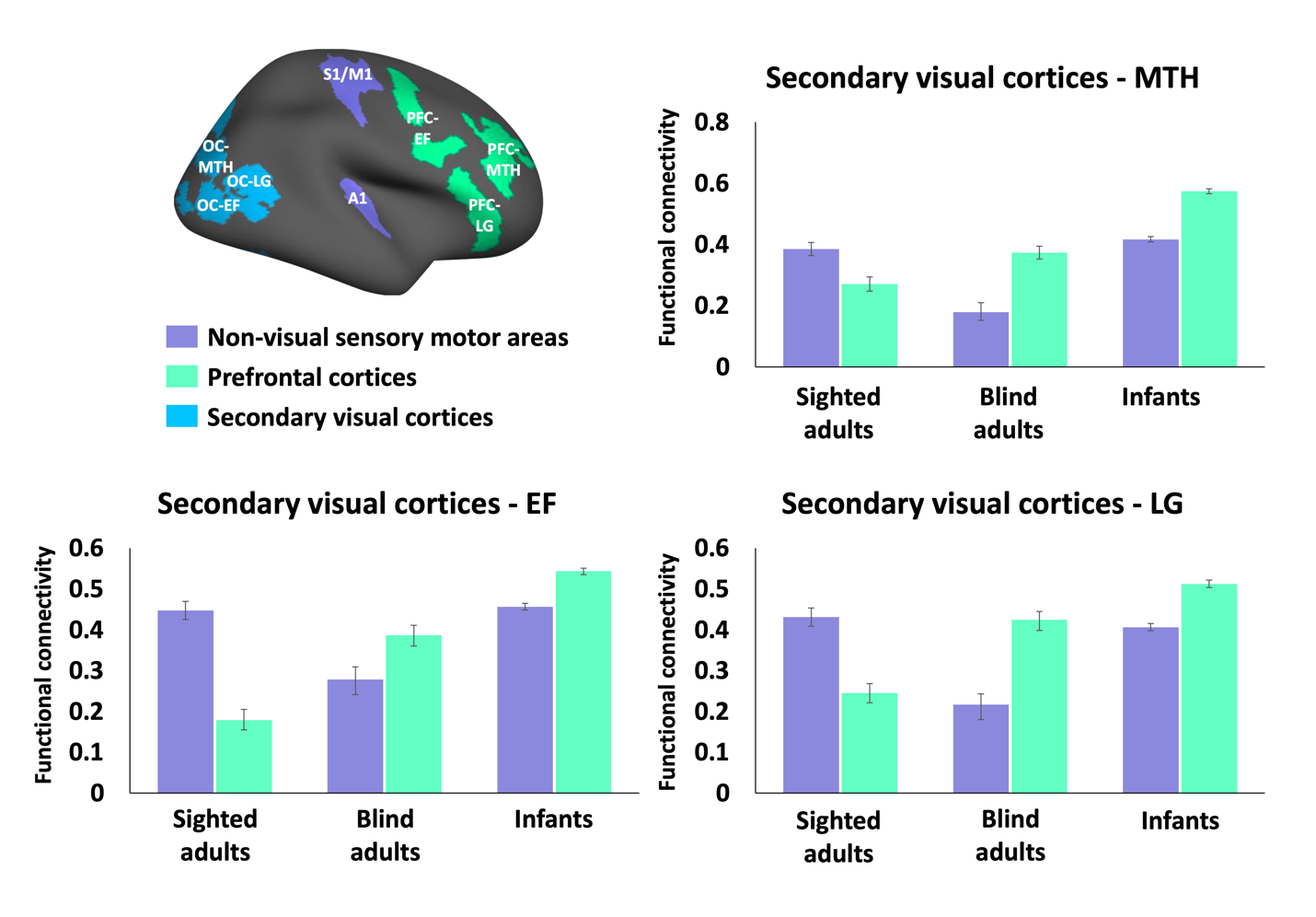
**Figure S2 Functional connectivity of three secondary visual regions.** The bar graph showed the resting state functional connectivity of three secondary visual regions to non-visual sensory-motor networks (purple) and prefrontal cortices (green) in sighted adults, blind adults and sighted infants, averaged across occipital, PFC and sensory-motor ROIs (A1 and S1/M1). Regions of interest displayed on the upper left. (PFC: prefrontal cortices; OC: occipital cortices; MTH: math-responsive region; LG: language-responsive region; EF: executive function-responsive (response-conflict) region.

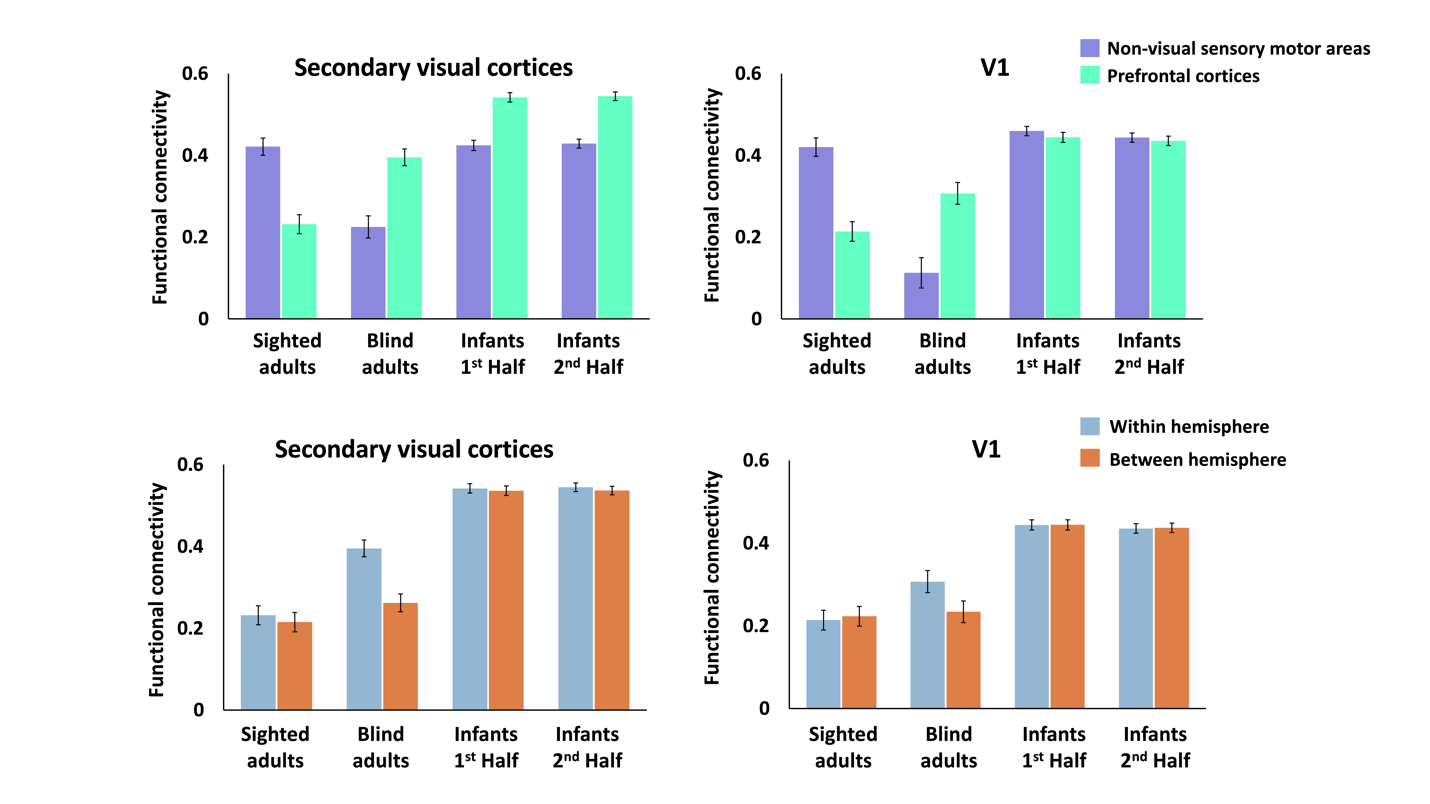

**Figure S3 Infants split-half results:** We split the infants dataset into two halves (n1 = 238, n2 = 237) and conducted split-half cross-validation. The functional connectivity between the secondary visual cortices (upper left) and V1 (upper right) to non-visual sensory-motor networks (purple) and prefrontal cortices (green) was shown in the upper row for sighted adults, blind adults, and two independent subgroups of sighted infants. The lower row displayed the functional connectivity within the hemisphere (blue) versus between hemispheres (orange) from the secondary visual areas (lower left) and V1 (lower right) to the prefrontal cortices.

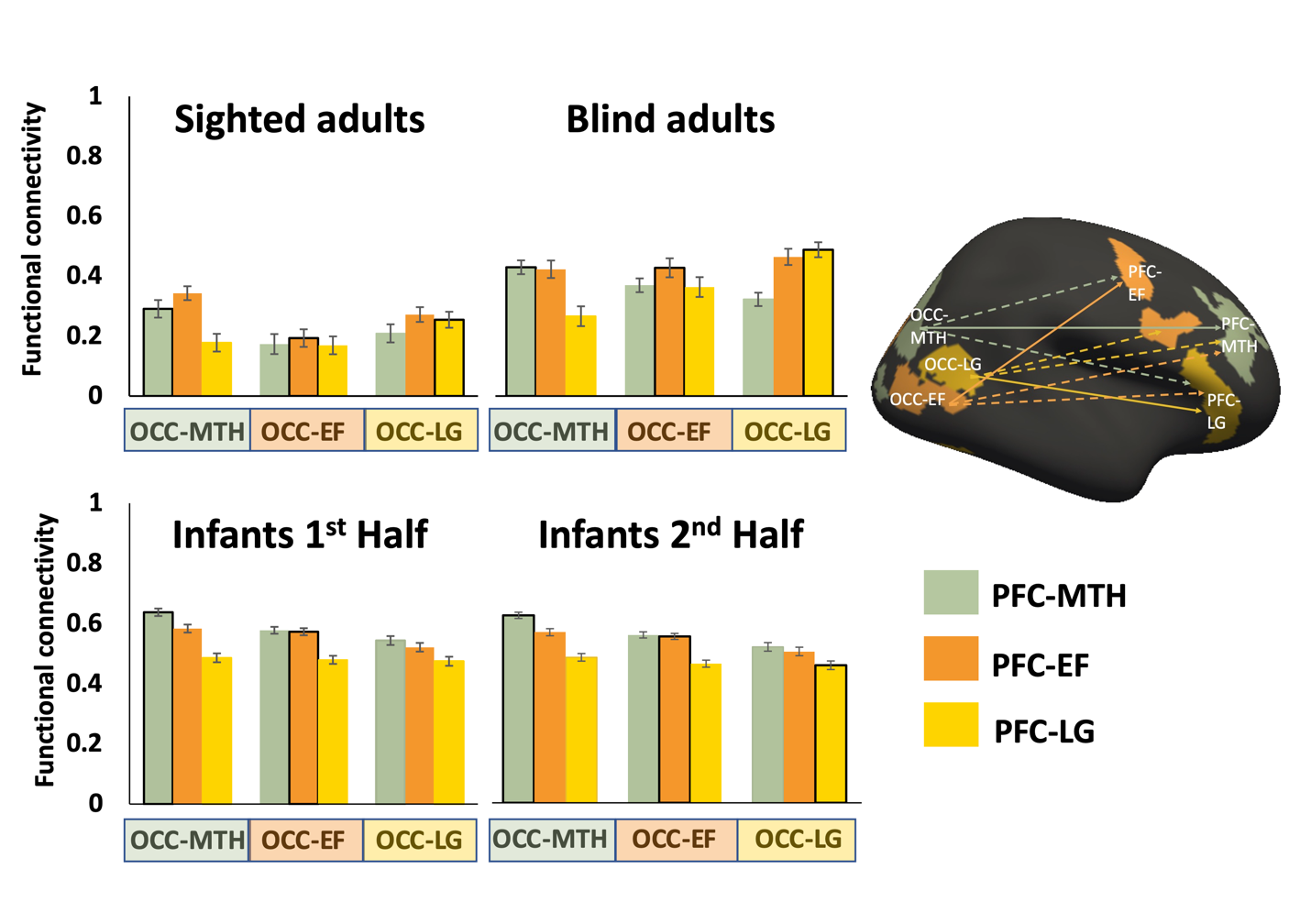

**Figure S4 Infants split-half results:** Occipito-frontal functional connectivity across different sub-regions of prefrontal (PFC) and occipital cortex (OCC) in sighted adults, blind adults and two independent subgroups of sighted infants (infants were randomly assigned to two subgroups, n1 = 238, n2 = 237). PFC: prefrontal cortices; MTH: math-responsive region; LG: language-responsive region; EF: executive function (response-conflict) region

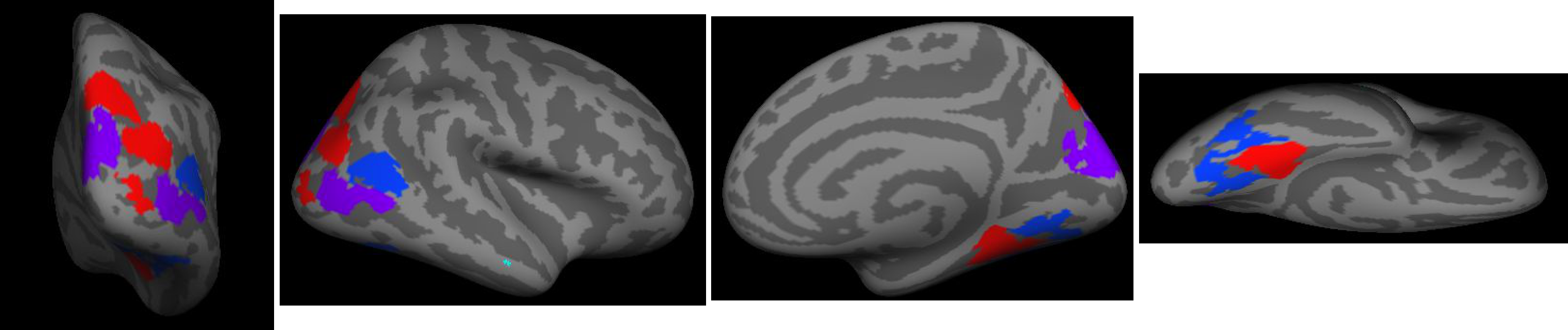

**Figure S5 Full views of the occipital ROIs.** Occipital math-responsive regions (red) were more active when solving math equations than comprehending sentences. Occipital language-responsive regions (blue) were more active when comprehending sentences than solving math equations; Occipital executive function (response-conflict) regions were more active during response inhibition (no-go) trials than active go trials during an auditory no-go task (Kanjlia et al., 2016, 2021; Lane et al., 2015). The occipital ROIs were defined based on group comparisons blind > sighted in a whole-cortex analysis. All three occipital ROIs were defined in the right hemisphere. Any overlapping voxels between ROIs were removed and not counted toward any ROIs.

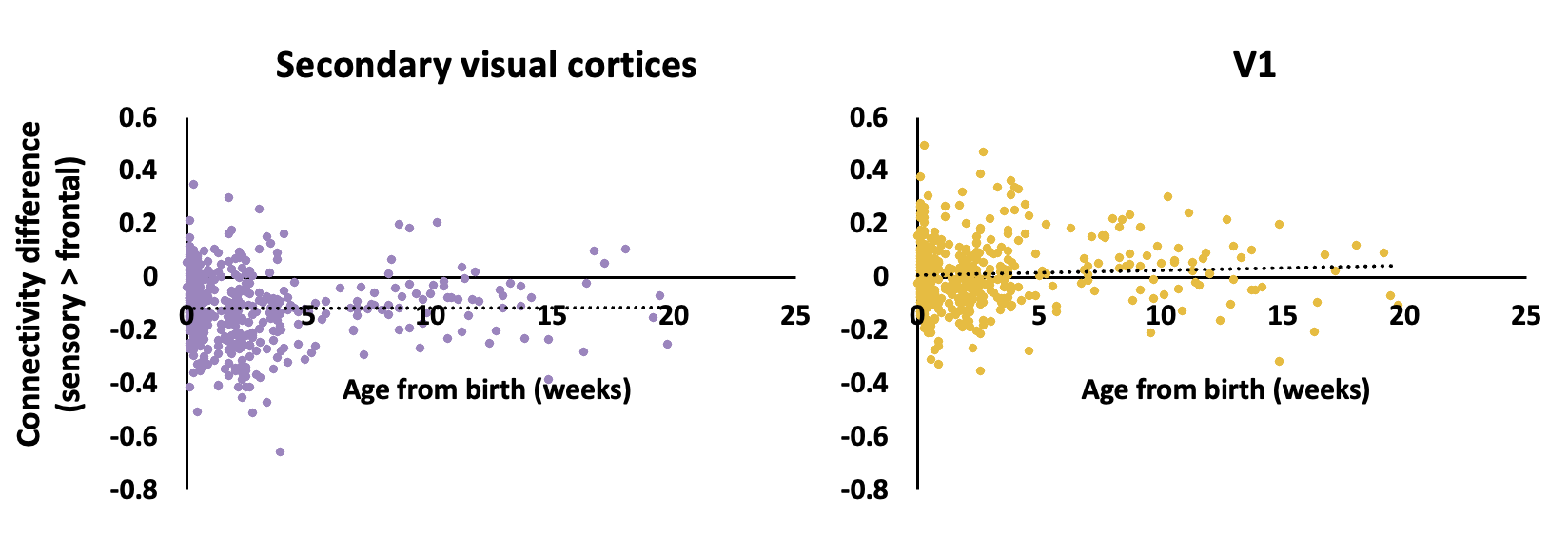

**Figure S6 The correlation of discrepancy in connectivity of visual cortex with age.** The scatter plot showed the correlation of discrepancy in connectivity of visual cortex (secondary visual cortices (left) and V1 (right)) to non-visual sensory areas and to prefrontal cortex with age after birth in infants. Data points represent individual participants.

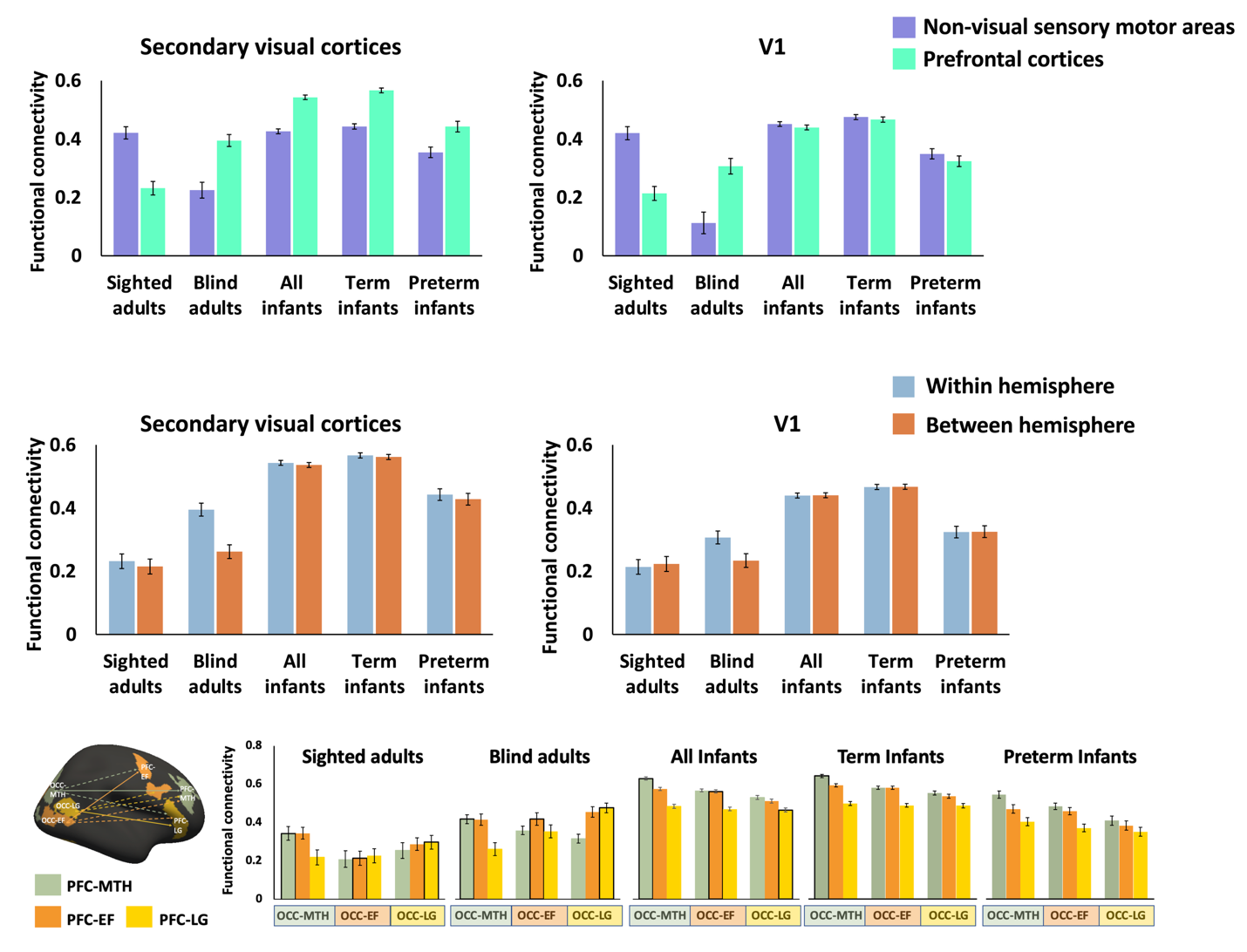

**Figure S7 Preterm and term infants results**: We also compared the results for preterm (*n* = 90) and term infants (*n* = 385) and found similar outcomes. The functional connectivity between the secondary visual cortices (upper left) and V1 (upper right) to non-visual sensory-motor networks (purple) and prefrontal cortices (green) is shown in the upper row. The middle row displays the functional connectivity within the hemisphere (blue) versus between hemispheres (orange) from the secondary visual areas (lower left) and V1 (lower right) to the prefrontal cortices. The lower row displays the occipito-frontal functional connectivity across different sub-regions of the prefrontal (PFC) and occipital cortex (OCC). MTH: math-responsive region; LG: language-responsive region; EF: executive function (response-conflict) region.

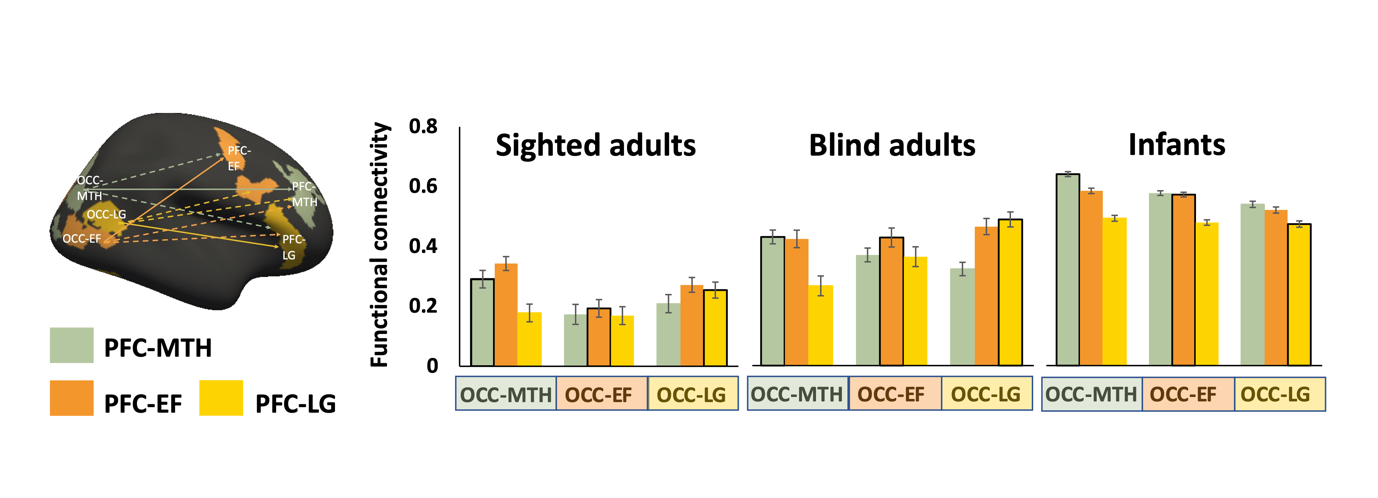

**Figure S8** **Occipito-frontal functional connectivity across different sub-regions of prefrontal (PFC) and occipital cortex (OCC) in sighted adults, blind adults, and sighted infants.** Sub-regions (regions of interest) were defined based on task-based responses in a separate dataset of sighted (frontal) and blind (frontal and occipital) adults (Kanjlia et al., 2016, 2021; Lane et al., 2015). PFC/OC-MTH math-responsive regions were more active when solving math equations than comprehending sentences. PFC/OC-LG language-responsive regions were more active when comprehending sentences than solving math equations; EF: executive function (response-conflict) regions were more active during response inhibition (no-go) trials than active go trials during an auditory no-go task (Kanjlia et al., 2016, 2021; Lane et al., 2015). In blind adults (top right) these regions show biases in connectivity related to their function i.e., language-responsive PFC is more correlated with language responsive OCC. No such pattern is observed in infants.

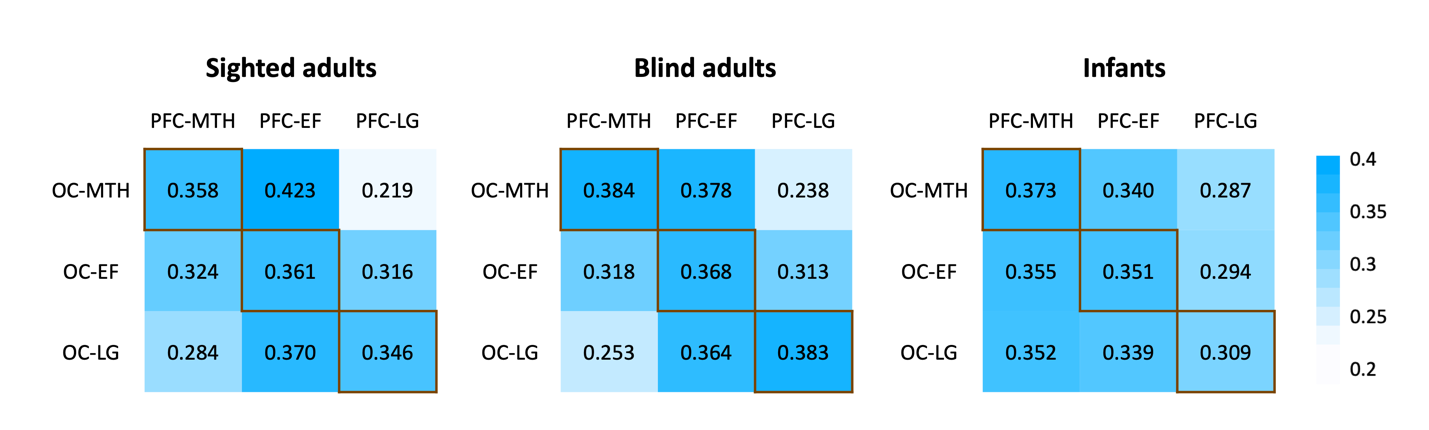

**Figure S9** The resting-state functional connectivity matrices between secondary visual areas to prefrontal regions in sighted adults, blind adults, and sighted infants. PFC: prefrontal cortices; OC: occipital cortices; MTH: math-responsive region; LG: language-responsive region; EF: executive function (response-conflict) region.

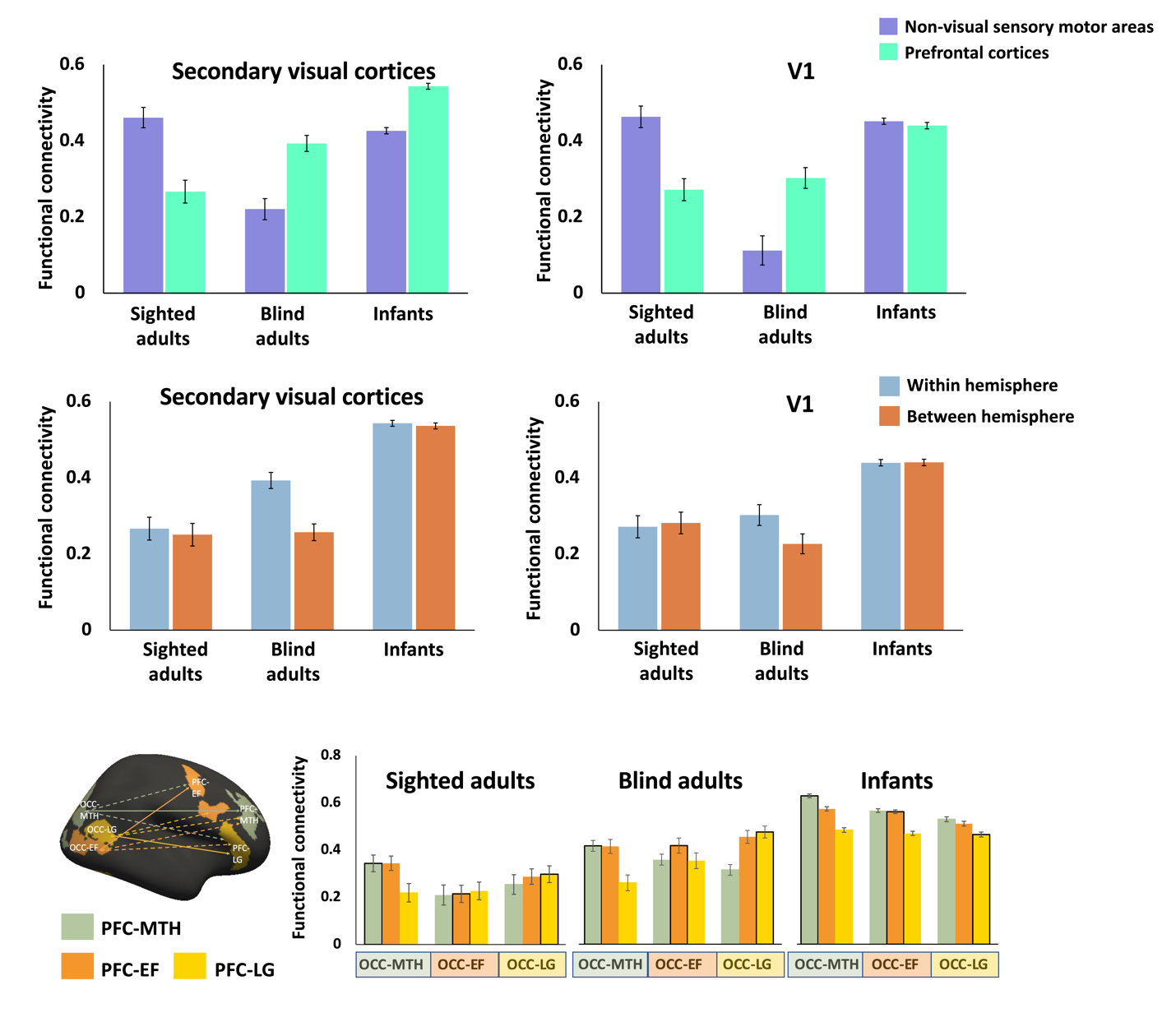

**Figure S10 Age-matched adults subgroup results**: We performed analyses in age-matched subgroups of sighted controls (n = 29, average age across scans: M = 43, SD = 13.04) and blind adults (n = 29, average age across scans: M = 43.24, SD = 15.75). The functional connectivity between the secondary visual cortices (upper left) and V1 (upper right) to non-visual sensory-motor networks (purple) and prefrontal cortices (green) is shown in the upper row. The middle row displays the functional connectivity within the hemisphere (blue) versus between hemispheres (orange) from the secondary visual areas (lower left) and V1 (lower right) to the prefrontal cortices. The lower row displays the occipito-frontal functional connectivity across different sub-regions of the prefrontal (PFC) and occipital cortex (OCC). MTH: math-responsive region; LG: language-responsive region; EF: executive function (response-conflict) region.

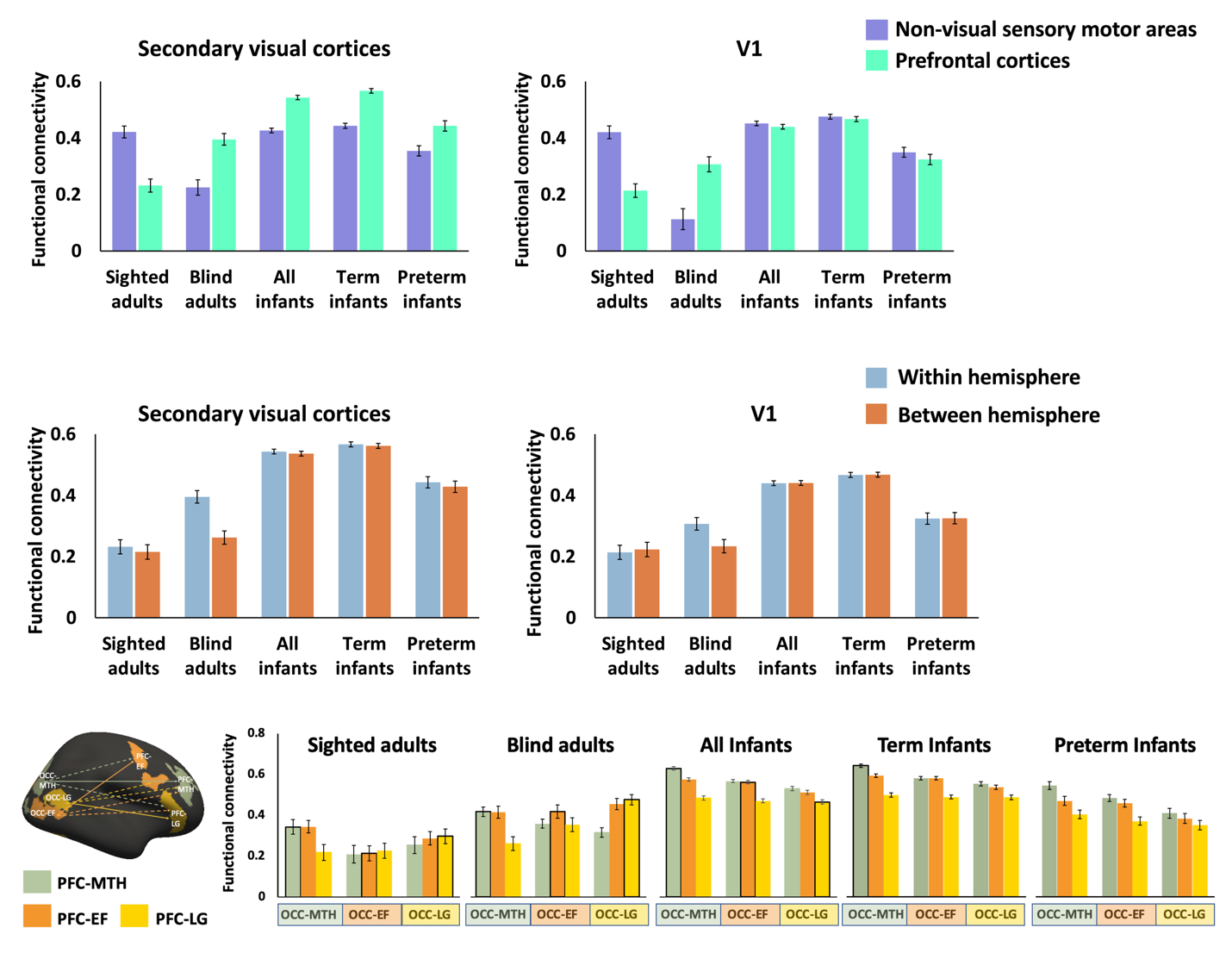

**Figure S11 Results of dataset excluding infants with radiology scores of 4 or 5**: We preform our analysis on the dataset that excluded the infantss who had a radiology score of 4 or 5, and found that the results remained the same. The functional connectivity between the secondary visual cortices (upper left) and V1 (upper right) to non-visual sensory-motor networks (purple) and prefrontal cortices (green) is shown in the upper row. The middle row displays the functional connectivity within the hemisphere (blue) versus between hemispheres (orange) from the secondary visual areas (lower left) and V1 (lower right) to the prefrontal cortices. The lower row displays the occipito-frontal functional connectivity across different sub-regions of the prefrontal (PFC) and occipital cortex (OCC). MTH: math-responsive region; LG: language-responsive region; EF: executive function (response-conflict) region.

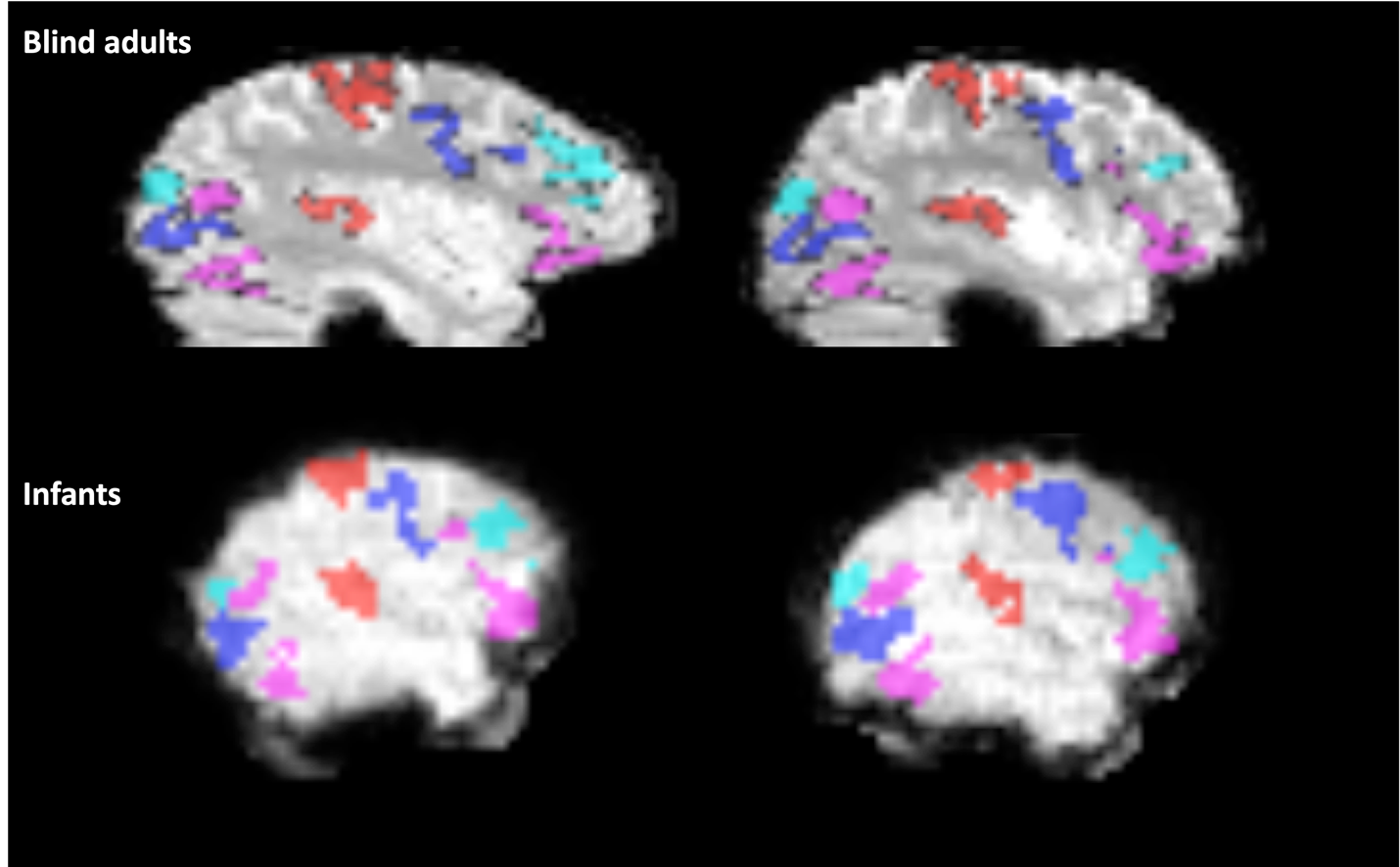

**Figure S12 Examples of ROI alignment on individual functional images.** The top row shows two blind adults, and the bottom row shows two infants. ROIs are color-coded as follows: A1 and S1/M1 in red, math-responsive frontal and occipital regions in cyan, language-responsive frontal and occipital regions in pink, and response-conflict responsive frontal and occipital regions in blue.

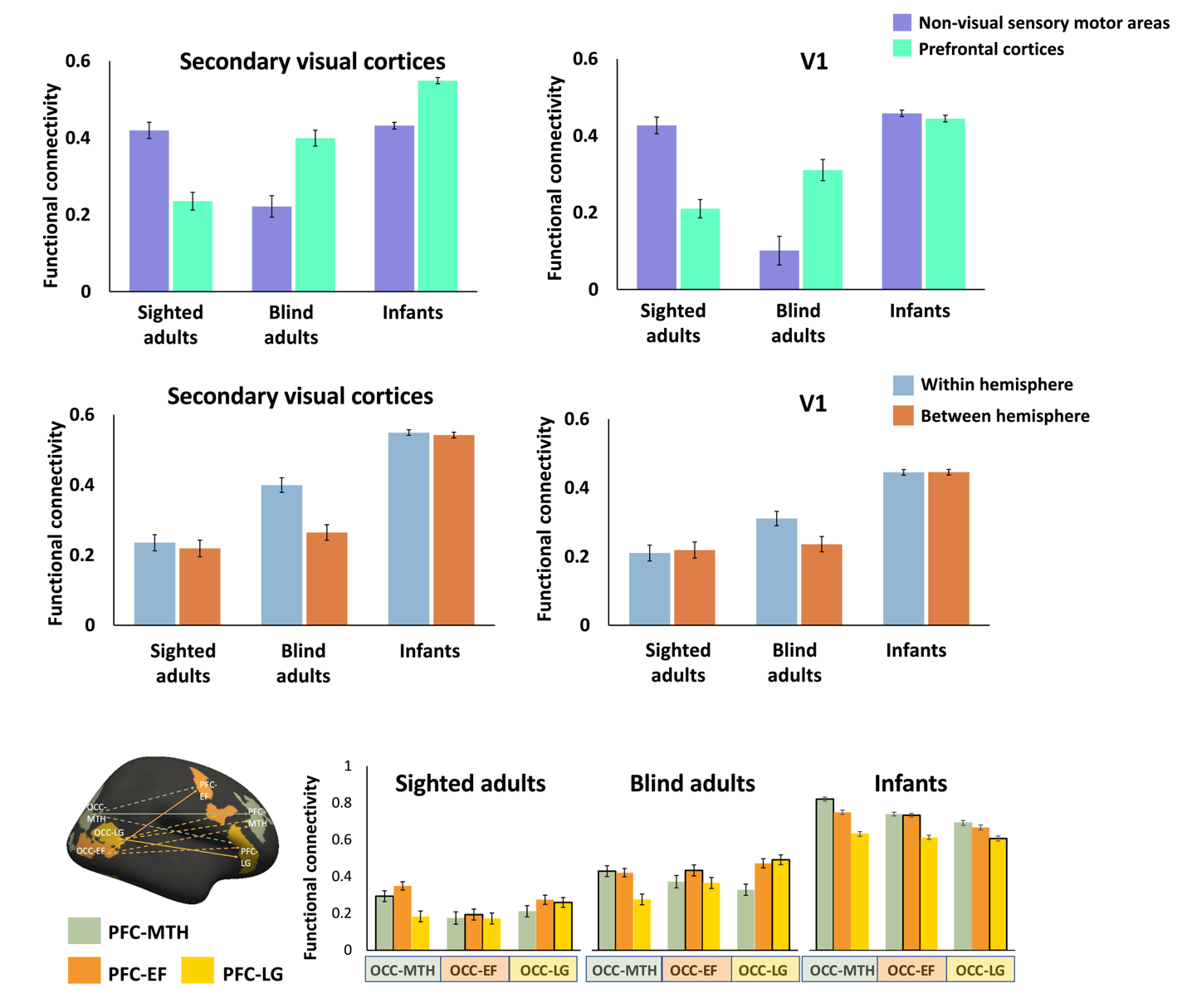

**Figure S13 Results of datasets excluding adults with signal dropout.**We performed our analysis on the datasets excluding the adult participants who showed signal dropout in one ROI (one sighted adult and two blind adults) and found that the results remained the same. The functional connectivity between the secondary visual cortices (upper left) and V1 (upper right) to non-visual sensory-motor networks (purple) and prefrontal cortices (green) is shown in the upper row. The middle row displays the functional connectivity within the hemisphere (blue) versus between hemispheres (orange) from the secondary visual areas (lower left) and V1 (lower right) to the prefrontal cortices. The lower row displays the occipito-frontal functional connectivity across different sub-regions of the prefrontal (PFC) and occipital cortex (OCC). MTH: math-responsive region; LG: language-responsive region; EF: executive function (response-conflict) region.

**Table S1** Post-hoc Bonferroni-corrected paired t-test for the connectivity between occipital regions to prefrontal regions in infants:

|  | Infants |
| --- | --- |
| OC-MTH to PFC-MTH vs. OC-MTH to PFC- EF | *t* _(474)_ = 10.7, *p* < 0.001 |
| OC-MTH to PFC-MTH vs. OC-MTH to PFC- LG | *t* _(474)_ = 24.438, *p* < 0.001 |
| OC-MTH to PFC- EF vs. OC-MTH to PFC- LG | *t* _(474)_ = 24.438, *p* < 0.001 |
| OC-LG to PFC-MTH vs. OC-LG to PFC- EF | *t* _(474)_ = 3.667, *p* < 0.001 |
| OC-LG to PFC-MTH vs. OC-LG to PFC-LG | *t* _(474)_ = 11.269, *p* < 0.001 |
| OC-LG to PFC-EF vs. OC-LG to PFC-LG | *t* _(474)_ = 7.581, *p* < 0.001 |
| OC-EF to PFC-MTH vs. OC-EF to PFC- EF | *t* _(474)_ = 0.981, *p* = 0.981 |
| OC- EF to PFC-MTH vs. OC- EF to PFC- LG | *t* _(474)_ = 16.238, *p* < 0.001 |
| OC- EF to PFC- EF vs. OC- EF to PFC- LG | *t* _(474)_ = 13.536, *p* < 0.001 |
| PFC: prefrontal cortices; OC: occipital cortices; MTH: math-responsive region; LG: language-responsive region; EF: executive function (response-conflict) region*.* | |

**Table S2 ROI-wise noise ceiling and One-Way ANOVA results across the three groups**

|  | S(n = 50) | | CB(n = 30) | | Infants(n = 475) | | F | p | partial 𝜂^2^ |
| --- | --- | --- | --- | --- | --- | --- | --- | --- | --- |
|  | M | SD | M | SD | M | SD |  |  |  |
| A1_lh | 0.922 | 0.041 | 0.904 | 0.055 | 0.938 | 0.048 | 9.034 | 0.000 | 0.032 |
| A1_rh | 0.915 | 0.052 | 0.908 | 0.061 | 0.936 | 0.046 | 8.787 | 0.000 | 0.031 |
| PFC-MTH_lh | 0.921 | 0.049 | 0.923 | 0.036 | 0.941 | 0.046 | 5.886 | 0.003 | 0.021 |
| PFC-MTH_rh | 0.920 | 0.049 | 0.921 | 0.038 | 0.942 | 0.042 | 9.076 | 0.000 | 0.032 |
| PFC-LG_lh | 0.923 | 0.050 | 0.920 | 0.045 | 0.937 | 0.048 | 3.645 | 0.027 | 0.013 |
| PFC-LG_rh | 0.908 | 0.099 | 0.897 | 0.080 | 0.941 | 0.039 | 19.469 | 0.000 | 0.066 |
| PFC-EF_lh | 0.913 | 0.057 | 0.902 | 0.047 | 0.936 | 0.050 | 10.487 | 0.000 | 0.037 |
| PFC-EF _rh | 0.920 | 0.039 | 0.893 | 0.088 | 0.935 | 0.051 | 10.307 | 0.000 | 0.036 |
| SMC_lh | 0.903 | 0.069 | 0.928 | 0.049 | 0.942 | 0.051 | 12.839 | 0.000 | 0.044 |
| SMC_rh | 0.910 | 0.072 | 0.933 | 0.040 | 0.941 | 0.048 | 8.768 | 0.000 | 0.031 |
| OC-MTH_lh | 0.945 | 0.038 | 0.931 | 0.036 | 0.940 | 0.051 | 0.688 | 0.503 | 0.002 |
| OC-MTH _rh | 0.937 | 0.067 | 0.937 | 0.045 | 0.943 | 0.048 | 0.438 | 0.645 | 0.002 |
| OC-EF_lh | 0.957 | 0.033 | 0.943 | 0.038 | 0.935 | 0.050 | 4.536 | 0.011 | 0.016 |
| OC-EF_rh | 0.962 | 0.024 | 0.924 | 0.084 | 0.938 | 0.047 | 6.985 | 0.001 | 0.025 |
| OC-LG_lh | 0.947 | 0.047 | 0.935 | 0.039 | 0.934 | 0.056 | 1.406 | 0.246 | 0.005 |
| OC-LG_rh | 0.950 | 0.038 | 0.926 | 0.056 | 0.935 | 0.057 | 2.172 | 0.115 | 0.008 |
| V1_lh | 0.953 | 0.027 | 0.941 | 0.047 | 0.921 | 0.064 | 6.831 | 0.001 | 0.024 |
| V1_rh | 0.949 | 0.039 | 0.943 | 0.046 | 0.922 | 0.065 | 5.340 | 0.005 | 0.019 |

PFC: prefrontal cortices; OC: occipital cortices; MTH: math-responsive region; LG: language-responsive region; EF: executive function (response-conflict) region*.*

**Results - The effects across the three different secondary visual regions to non-visual sensory areas and prefrontal regions in infants**

Secondary visual regions (math, language, response-conflict) by ROIs (PFC, non-visual sensory) repeated measure ANOVA were conducted in infants. A significant interaction effect was found between the visual regions and ROIs (*F*_(2, 948)_ = 136.968, *p* < 0.001). A post-hoc Bonferroni-corrected paired t-test revealed a similar connectivity pattern across the three secondary visual regions, which exhibited stronger connectivity to prefrontal regions than non-visual sensory regions. However, the largest mean difference was observed in the occipital math-responsive region, followed by the language-responsive region, with the smallest difference found in the occipital conflict-responsive region (connectivity to non-visual sensory and to PFC, occipital math: mean difference: -0.209, *t* _(474)_ = -24.546, *p* < 0.001; occipital language: mean difference: -0.141, *t* _(474)_ = -16.674, *p* < 0.001; occipital conflict: mean difference: -0.114, *t* _(474)_ = -13.755, *p* < 0.001).
